# HArtMuT - Modeling eye and muscle contributors in neuroelectric imaging

**DOI:** 10.1101/2022.08.19.504507

**Authors:** Nils Harmening, Marius Klug, Klaus Gramann, Daniel Miklody

## Abstract

Magneto- and electroencephalography (M/EEG) measurements record a mix of signals from the brain, eyes, and muscles. These signals can be disentangled for artifact cleaning e.g. using spatial filtering techniques. However, correctly localizing and identifying these components relies on head models that so far only take brain sources into account. We thus developed the Head Artefact Model using Tripoles (HArtMuT). This volume conduction head model extends to the neck and includes brain sources as well as sources representing eyes and muscles that can be modeled as single dipoles, symmetrical dipoles, and tripoles.

We compared a HArtMuT 4-layer Boundary Element Model (BEM) with the EEGLAB standard head model on their localization accuracy and residual variance (RV) using a HArtMuT Finite Element Model as ground truth. We also evaluated the RV on real-world data of mobile participants, comparing different HArtMuT BEM types with the EEGLAB standard head model. We found that HArtMuT improves localization for all sources, especially non-brain, and localization error and RV of non-brain sources were in the same range as those of brain sources. The best results were achieved by using cortical dipoles, muscular tripoles, and ocular symmetric dipoles, but dipolar sources alone can already lead to convincing results.

We conclude that HArtMuT is well suited for modeling eye and muscle contributions to the M/EEG signal. It can be used to localize sources and to identify brain, eye, and muscle components. HArtMuT is freely available and can be integrated into standard software.

## Introduction

Magnetoencephalography (MEG) and Electroencephalography (EEG) are methods to non-invasively measure the electrical activity of populations of cortical neurons in humans and other species. EEG provides high temporal but only limited spatial resolution, because of the sensors being placed on the scalp outside the low-conductive skull. Due to volume conduction, only a spatially smeared linear mixture of all cortical sources, that are active in parallel, arrives at the sensor level [Haufe et al., 2014]. MEG has better spatial resolution than EEG, since the magnetic fields are less distorted by the mixture of different conductive tissues (brain, cerebrospinal fluid (CSF), skull, scalp), but suffers from low signal-to-noise ratio in particular for deeper sources. In addition, MEG is sensitive primarily to tangential-oriented sources and is more demanding in its recording procedure.

Both MEG and EEG, referred to as M/EEG in the following, record volume conducted signals composed of cortical sources that are active in parallel and mixing in a linear fashion. Dissociating the different sources can be solved by source separation techniques [Pearson, 1901, Makeig et al., 1996a, Blankertz et al., 2011], that introduce a number of assumptions to solve the ill-posed inverse problem [Sarvas, 1987]. Mathematically dissociating cortical sources in M/EEG signals is further complicated by other sources contributing to the recorded signal, like physiological non-brain sources (eye movement, muscle activity) and mechanical (cable sway in EEG) or electrical sources (noise stemming from stimulation monitors and other devices).

While mechanical and electrical sources often show invariant patterns, that can be easily dissociated, eye movement and muscle activity of facial and neck muscles are often less easy to identify. Importantly, these classes of non-brain sources contribute significantly to the signal as they are located outside the skull and show higher power compared to the minuscule EEG signal. Signals from eye movements and blinks can be orders of magnitude larger than brain-generated electrical potentials. Therefore, activity stemming from eye movements, the closure of the lid, and its necessary muscular activity together with neck and face muscle activity are the most common types of artefacts in EEG recordings [Joyce et al., 2004, Croft and Barry, 2000].

This kind of non-brain activity is traditionally considered to be artefactual, even though eye movement and facial mimicry significantly contribute to or are a result of cognitive and affective processes [König et al., 2016]. As a consequence, a number of artefact rejection methods have evolved to identify and remove such kind of activity from the recorded signal. Rejecting contaminated trials, however, causes substantial data loss, and restricting eye movements and blinks might limit the experimental design and could impact the cognitive processes under investigation. Thus, newer algorithms attempt to remove artefact specific features from the signal without deleting contaminated time periods to preserve as much data as possible [Dimigen and Ehinger, 2021, Jung et al., 2000].

Among these algorithms are blind source separation (BSS) techniques [Urigüen and Zapirain, 2015] that decompose the signal mostly based on its spatio-temporal statistics into different sources after which neural sources are identified and separated. The most commonly used approach is Independent Component Analysis (ICA) which separates the sources based on the principle of maximal statistical independence [Makeig et al., 1996b]. The assumption that neural and artefactual sources are independent implies the possibility of sorting them into sources of different origins. This can be done manually by an expert or by applying automatic routines [Winkler et al., 2011, Pion-Tonachini et al., 2019].

Eventually, the sources that have been identified as originating in the brain, are further localized using source localization approaches [Oostendorp and Van Oosterom, 1989, Oostenveld et al., 2011, Mosher et al., 1992]. Finding a source in general requires computing forward solutions by a head model, so-called leadfields. Their difference from the original data is subject to minimization, where the exact source position is the optimization parameter. The same approach to localize brain sources could thus be used to also localize other physiological sources. However, to our knowledge, no head model actually provides leadfields that incorporate the eyes and face and neck muscles in the forward solution.

Furthermore, the simulation of different signal sources including those of eyes and muscles is useful for educational and validation purposes, as it has been done for neural origins [Krol et al., 2018]. This also involves a biophysical forward model, that mimics the field propagation within the head, and has so far been done almost exclusively for neural sources in existing models.

In order to improve the source localization accuracy, to offer a better guide for users to distinguish brain from non-brain sources and to provide a model, which is suitable for eye and muscle simulations, we developed the Head Artefact Model using Tripoles (HArtMuT), incorporating eyes, facial muscles and muscles in the dorsal neck region into EEG leadfields.

In this study, exact source positions and orientations were extracted from the open-source, high-resolution Multimodal Imaging-Based Detailed Anatomical (MIDA) model [Iacono et al., 2015]. Dipolar, tripolar and symmetric dipolar structures for realistic artefact modeling were physiologically motivated and placed onto these positions. A 4-shell Boundary Element Method (BEM) head model of the anatomy of Colin27 [Holmes et al., 1998] with neck extended scalp mesh was used to evaluate the artefact model on both simulated HD-FEM data and ICA patterns from EEG recordings.

The final HArtMuT is publicly available^1^ as Boundary and Finite Element Method (BEM/FEM) model for different commonly used standard head anatomies and is ready to be implemented into standard approaches of all kinds.

## Source modeling

In the lower frequencies (< 1 kHz), the basis of common electrical models of volume conduction in the human head with conductivity *σ* is the quasi-electrostatic assumption [Häamäläinen et al., 1993], which leads to a Poisson equation that has to be solved in order to model the field distributions (see e.g. Sarvas [1987]):

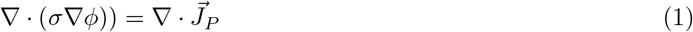

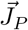 are the sources of primary currents that lead to the volume conduction *σ*∇*ϕ*, hence the electrical potential *φ*. 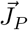 are usually modeled to be evoked by a superposition of point sources like single dipoles called the equivalent current dipole (ECD). The sources produce a local current flow which leads to a return current in all connected conductive tissues. The infinite homogeneous solutions of eqn. 1 for a given geometry with the ECD as *J_P_* are commonly used to simulate the electrical potential of a firing population of neurons. These solutions are unique and can be numerically determined. The most common numerical methods nowadays are Boundary Element Method (BEM), Finite Element Method (FEM), and Finite Differences Method (FDM). In general, FEM/FDMs can incorporate more detail in anatomy and conductivity, such as anisotropy or certain local inhomogeneities, but are computationally much more expensive.

In the following, the physiology of the different source types, their electrical fields, and corresponding source models are examined in detail, demonstrating that the currents of dipolar 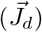 and tripolar 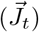 source models are reasonable approximations for 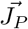, when modeling EEG contributors.

### Cortical sources

The main neuronal activity is believed to be based on membrane potentials and electrical currents through ions. Whether being Action Potentials (AP) along the axons or Postsynaptic Potentials (PSP), they produce a current flow through the membrane, that leads to a secondary return current flow through the neuron’s surrounding, which can be measured outside the head [Buzsáki et al., 2012]. In particular scalp EEG is assumed to be primarily elicited by the mean field produced by PSPs of larger assemblies of neocortex layer pyramidal cells. The ECD is a widely accepted approximation for the electric field of such a population of parallelly aligned and synchronously firing neurons. For a detailed derivation of the ECD from the contribution of principal neocortical neurons to the EEG, please refer to Murakami and Okada [2006].

The far-field of an ECD is described by an analytic expression, which is then used in combination with the head model to describe the resulting volume conduction within the whole head. In Cartesian coordinates, this electrical far-field potential of a dipolar current source in an infinite homogeneous conductive medium is

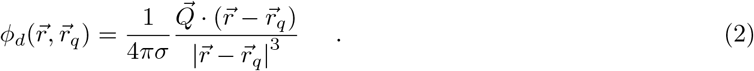

And the current density is calculated as

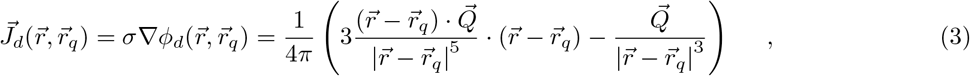

where 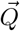 is the local current density produced by the source, 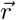 the observed point, and 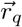 the position of the source.

### Ocular sources

Ocular artefacts in the EEG are caused by eye movements or blinks and are associated with higher amplitudes and lower frequencies than those of cortical EEG signals. The underlying sources are mainly permanent charge distributions in the eyeballs, but additionally include the eye muscles, that produce an electrical field, when active, and the closure of the eyelids, which modifies the volume conduction. The eyes are frequently the components of strongest amplitudes and can be easily identified by their specific frontal patterns. Eye movements of both eyes are generally linked in healthy humans which leads to a joint field produced by both eyes equally. Since the potentials are correlated among both eyes, single eye sources cannot be separated by ICA/PCA. Hence, this symmetry has to be incorporated into the model, which is taken care of by investigating both single and symmetric eye models.

### Physiology

The eyes are electrically charged at different structures as depicted in fig. 1. Firstly, the photoreceptors in the retina maintain a standing negative resting potential in order to turn incoming light into an excitation. In the pigment epithelium, this results in an electrical signal, which then propagates through the optic nerve to the visual cortex. Contrary to this strong negative charge distribution, fewer positive potassium and sodium ions are located in the outer segment of the photoreceptor cells, forming the overall membrane potential of around –40mV. When stimulated, the photoreceptors become even more hyperpolarized. Opposing the retina’s negativity is the overall positivity of the cornea [Iwasaki et al., 2005]. In its function to protect the frontal eye from physical and chemical agents, it transports positively charged sodium ions inward and negatively charged chloride ions outward. Therefore the surface, the corneal epithelium, is positively charged, as opposed to less positivity inside. During eye movements or blinks, the described charge distributions move and their electrical field changes. Since it is these changes, that are recorded by EEG [Berg and Scherg, 1991], an equivalent ocular model is expected to describe the charge distribution difference during movement rather than the static charge distributions. Additionally, the eyeball rotation is driven by small muscles around the eyes, that produce an electrical field when active. The closure of the eyelid is thought to additionally change the local field by altering the geometry of the volume conductor [Plöchl et al., 2012].

**Figure 1:**
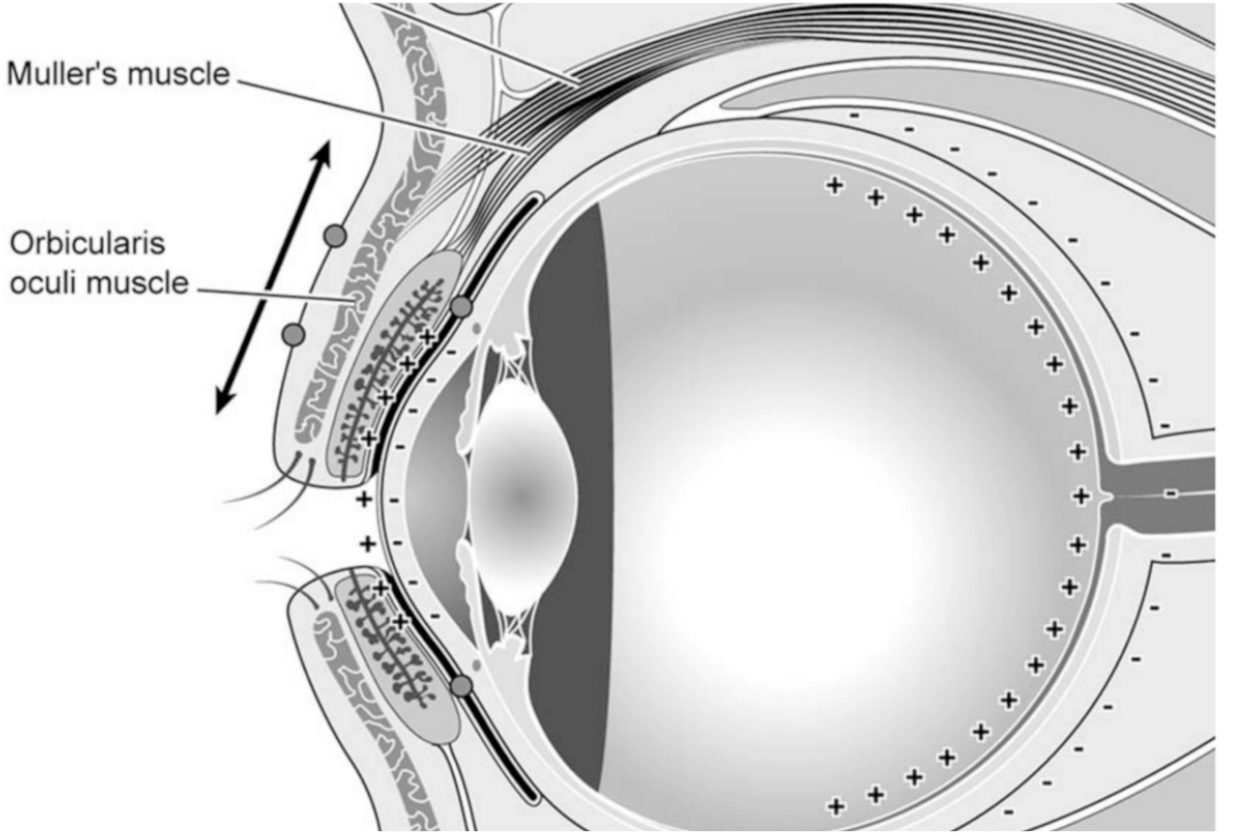
Anatomy and charge distribution of the eye and eyelid. On the right, the negatively charged (—) retina and its opposing smaller positive charges (+), and on the left, the inside negativity of the cornea and its surface positivity. An eyeball rotation results in charge distribution differences, that are visible in the EEG. The eyelid itself consists of the Orbicularis oculi muscle and its sliding is functioned by Muller’s muscle. The eyeball rotation muscles are omitted in this schematic depiction for the sake of simplicity. Reprinted from Iwasaki et al. [2005], Copyright 2004, with permission from Elsevier.

### Previous approaches to model eye activity

Some studies already investigated modeling the contribution of eye activity to EEG signals, where the strong corneo-retinal potential was mostly approximated by one equivalent ocular dipole with its negative pole posteriorly (directed towards the retina) and its positive pole anteriorly (directed towards the cornea). Using Electrooculography (EOG) recorded signals during vertical and horizontal eye movements and eye blinks, Berg and Scherg [1991] observed, that ‘a *reasonable fit was only obtained if the equivalent dipoles were allowed to take up different locations and orientations depending on the type of eye activity’.* A single dipole was determined for each movement across subjects. However, the difference dipole of a single eye movement was assumed to be approximately stationary due to asking the subjects to move their eyes always by the same fixed angle. Therefore, one might conjecture, that even more equivalent dipoles for the same movement direction but different angles are needed. One specific dipole per eye was also determined to approximate the EOG signal during eye blinks. This dipole bears analogy to that of vertical eye movements, which reflects Bell’s phenomenon: *‘Spontaneous blinks produced small eye movements directed down and inward, whereas slow or forced blinks were associated with delayed upward eye rotations*’, as stated by [Iwasaki et al., 2005]. They undertook a detailed EOG analysis of eye and lid movements, reasoning that *‘EOG signals during vertical eye- and lid-movements are greatly influenced by the eyelids’,* since a closed eyelid conducts the ocular dipole field to the frontopolar scalp region within the scalp tissue.

### Conclusions regarding modeling eye activity

In order to mirror the actual biological charge distribution differences correctly, dipolar sources are placed into retina and cornea, respectively. Accompanied by eyeball rotations, activity of the extraocular muscles is expected, as well as Muller’s muscle activity due to eyelid closures during blinks. This muscular activity is included in the muscular model.

Additionally, we hypothesize that both eyes move simultaneously and synchronously during regular usage in healthy subjects, which leads to one electric field and therefore patterns stemming from the superposition of two sources - one in each eye. To this end, we included symmetric dipolar fields into HArtMuT, that consist of a summation of leadfields of left and right eye positions with identical orientation vectors. The performance of the different ocular models (single and eye-symmetric dipoles together with tripolar sources in the eye muscles) was investigated. The same dipolar model as used for the brain (eq. (2) and (3)) and the same tripolar model as used for the muscles (eq. (6) and (7)) are obeyed, respectively. The modeling of the eyelid closure is incorporated by the inclusion of the eyeball meshes into the scalp mesh, which in turn, however, excludes the open eyelid state. Moreover, this approach lacks modeling eyeballs (and muscles) as their own tissue(s) with different conductivity than the remaining scalp.

### Muscular sources

Muscular activity is usually measured using bipolar electromyography (EMG) recordings. Several muscles are located in the face, on the human skull surface, and in the posterior and anterior neck regions. The relative strength of their activity patterns on the scalp surface is related to their location outside of the poorly conductive skull. This is why EMG activity is often observed in EEG recordings, although participants are restrained in their movements in traditional M/EEG protocols. Supracranial and facial muscles contribute to the EEG signal mainly in the higher frequency range starting at around 20 Hz. However, newer mobile EEG and Mobile Brain/Body Imaging studies [Jungnickel and Gramann, 2016, Gramann et al., 2021] demonstrated a strong contribution of neck muscles in moving participants with a contribution of lower frequencies in addition to the known higher frequency range from other muscles. Thus, it is even more important to dissociate this non-brain activity from the activity of interest, also in lower frequencies. Most of these muscle artefacts are larger in amplitude than the neuro-electric activity measured at the scalp. Their spatial scalp patterns differ from those of neural origin, since their sources are located directly in the scalp in close proximity to the EEG cap, and their field is thus more local. Additionally, due to their physiological properties, they have a rather tripolar field distribution, which has an impact on the patterns as well.

### Physiology

Skeletal muscles are composed of bundles of muscle fibers, that are long cylindrical cells containing many myofibrils consisting themselves of contractile units called sarcomeres. The muscle fibers and the corresponding motor neuron of the nervous system are connected via nerve fibers that divide into many terminal branches. Each chemical synapse of these axon terminals builds a so-called neuromuscular junction, ending on a specific region of the muscle fiber, called the motor end plate. When the nerve impulse for muscle contraction in form of an action potential reaches this neuromuscular junction, synaptic transmission to the muscle fiber begins. The action potential triggers a chemical process which leads to a depolarization of the resting membrane potential of approximately −70 mV. If the threshold potential (often between around –55 and –50 mV) is exceeded, the electrical impulse of the end plate’s potential travels down the traverse tubules (T tubules) into the interior of the muscle fiber and finally triggers the contraction of all sarcomeres [Merletti and Farina, 2016]. This depolarization and the subsequent re-polarization phase, called hyperpolarization, is followed by the afterhyperpolarization (AHP), in which the membrane potential falls below the resting potential until it returns to its normal resting voltage [Merletti and Farina, 2016].

The muscle fibers innervated by a single motor neuron are known as the muscle unit. Those muscle units innervated by the same motor neuron form together with its motor neuron a single motor unit (MU), such that a motor unit action potential (MUAP) is produced by the summation of the accompanying muscle unit’s end plate potentials. Upon muscle contraction, the generated MUAPs and their propagation from the muscle end plate to the tendons are recorded during electromyography (EMG) [Winter et al., 1994] and as unwanted artefacts during EEG. The propagation in several muscle fibers leads to a recorded signal, which is originating from multiple distributed source locations, and thus rather a mean field signal.

### Previous approaches to model muscle activity

While modeling ocular contributors to the EEG have already been discussed among researchers, muscular sources, to our knowledge, have not been added to any head model so far.

However, approaches in modeling EMG sources exist in particular for sources in the extremities. Some researchers used a simple dipole to approximate the MUAP’s curve [Boyd et al., 1978] [Winter et al., 1994], while others used (two) balanced tripoles pointing opposingly into the direction of the muscle fiber ends [Farina and Rainoldi, 1999], [Griep et al., 1982], [Merletti et al., 1999], [Roeleveld et al., 1997]. This is because the potential curve appears more tripolar with a skewness related to the tripolar asymmetry in propagation direction, when additionally taking the AHP into account, i.e. the negative sink after the action potential peak (fig. 2). As shown by Merletti and Farina [2016], the field of an equivalent tripolar source model has a similar curvature to the more realistic analytic waveform, that was originally developed by Rosenfalck [1969] and later modified by Nandedkar and Stålberg [1983].

**Figure 2:**
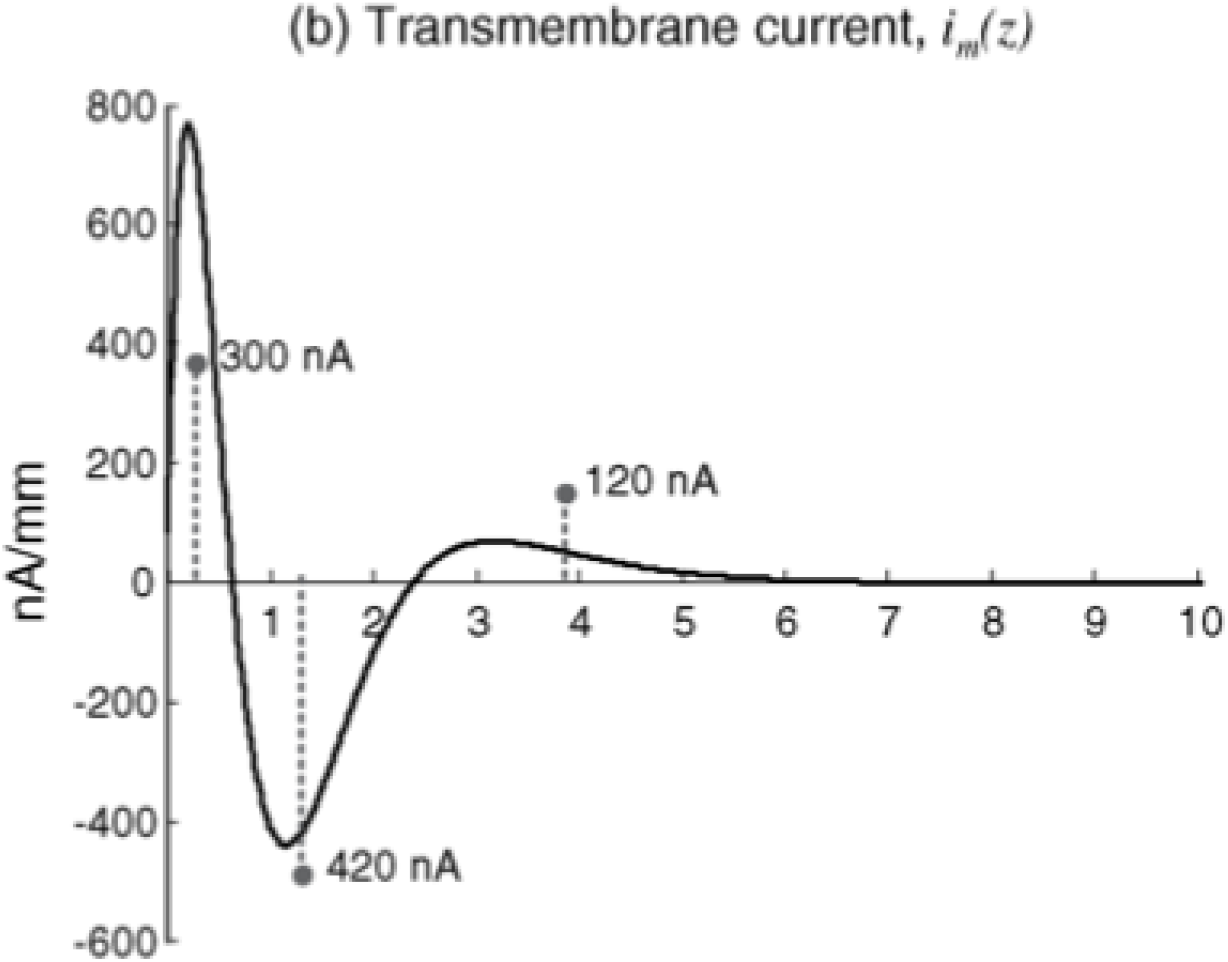
Simulated transmembrane current per unit length (based on Nandedkar and Stålberg [1983]) and the equivalent tripole source model as approximated by Merletti and Farina [2016]. The electric field of the first positive (300 nA) and the negative (420 nA) sources corresponds to the depolarization and the subsequent re-polarization phase. This is covered by an equivalent dipolar current source model. Adding the second positive charge (120 nA) a bit apart additionally incorporates the following small AHP bump. This constitutes the equivalent tripolar current source model *Tripole A*.

Used MUAP models vary in charge distribution and locations of the three monopoles, that form the tripole. Furthermore, the line, on which the three monopoles are placed, is not necessarily straight but can also be curved. All this has effects on the appearance of their surface potentials.

Fig. reprinted with permission from Merletti and Farina [2016]. Copyright 2016 by The Institute of Electrical and Electronics Engineers, Inc.

### Thoughts on MUAPs in the human head

In the human head, these muscular sources are found within what is usually modeled as scalp: a thinner sheet of conductive tissue bordering the non-conductive air and the much less conductive skull (the estimated conductivity ratio ranges from 1:40 to 1:80 [Goncalves et al., 2003, Clerc et al., 2005]). These rather sudden changes of conductivity also have an effect on the inner distributions of electrical potential and current flow. In fig. 3 one can observe that depending on the orientation of the tripole relative to the measured surface potential, it can have one, two, or three focal patterns on the surface. Moreover, the central (in our case positive) peaks of tripoles decay faster with distance than dipolar peaks in amplitude. Compared to brain sources, however, the missing spatial smearing effect of the CSF and skull additionally leads to more focal patterns of these scalp sources.

**Figure 3:**
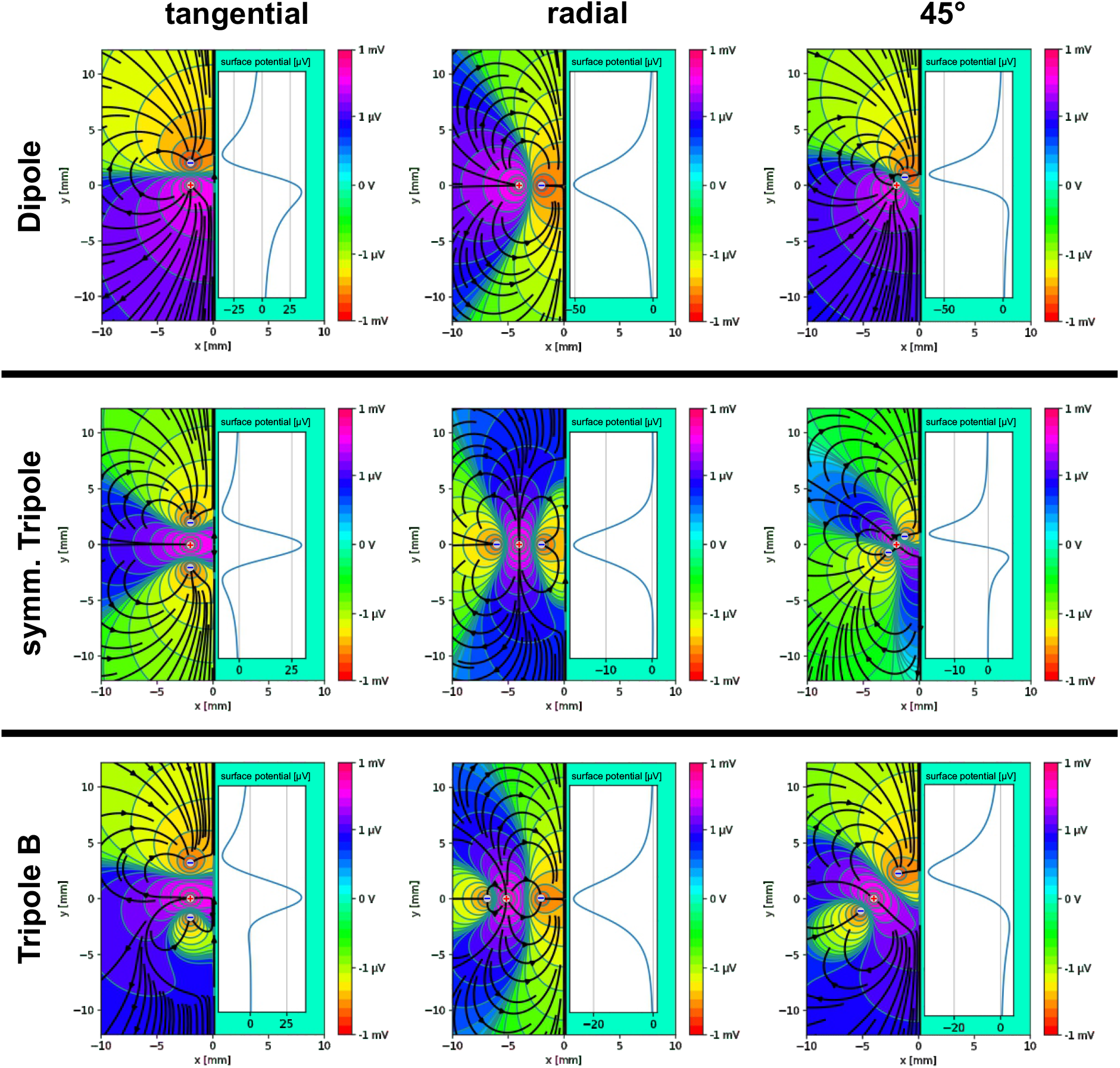
Comparison of scalp potential behavior for different source model types (dipole, sym. tripole, *Tripole B)* with three different orientations (tangential, radial, 45°) at similar source positions. Different orientations lead to different surface potential patterns: A tripole can look similar to a dipole or a monopole depending on its orientation, however, certain characteristics like the fall-off with distance differ. The dipole (upper row) consists of a double negative sink and a double positive source. A symmetric tripole (middle row) consists of two negative sinks at the same distance in opposite directions apart from a double positive source. *Tripole B* additionally has different distances leading to a rather dipolar potential distribution. The surface at *x* = 0 is to a non-conductive domain for *x >* 0, which simulates the scalp surface touching air. The lines depict different voltage levels (equipotential lines), while the flow lines represent the current flow (the density of the lines is arbitrary).

If tripoles are modeled similar to in Merletti and Farina [2016] (see fig. 2), the resulting field is in transition between dipole and tripole as the closer and stronger pair of sources (300nA/420nA) dominates the field.

### Conclusions regarding modeling muscle activity

Due to existing evidence for both dipolar and tripolar approaches, we decided to test both.

Noteworthy, little is known about EMG signals fall-off in current flow power within the muscle fibers [Kuiken et al., 2001], that, strictly speaking, one would additionally need to take into account. Since the potential form of EMG signals at different fiber positions is not simply a delayed versions of each other [Merletti and Farina, 2016], we place several myogenic sources along the signal’s two propagation directions from the neuromuscular junction (NMJ) to the tendons in the model.

Because the distance between the sources is in a similar range as the distance to the surface, a far field approximation, such as the ECDs used for neurons inside the brain, does not hold for sources outside the skull. Instead, the optimal model would be to directly model the current flow out of the muscle along its fibers using analytic solutions comparable to Nandedkar and Stålberg [1983]. However, such an implementation in existing head model software would not allow the usage of most existing inverse fitting techniques, that optimize location and orientation of point sources. We, therefore, model the tripolar structure as three single monopolar sources, as described by Merletti and Farina [2016]. In Cartesian coordinates, the electrical field potential of a single (monopolar) current source *φ_m_* in an infinite homogeneous conductive medium is calculated as

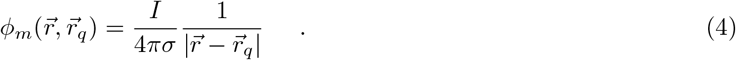

The corresponding current density 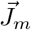 is given by

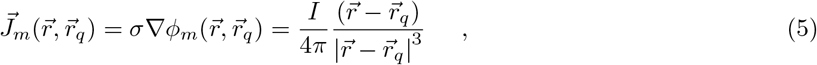

where *I* is the total current of the source. The field is radial symmetric.

In order to model the field of a tripole, three monopolar sources with different parameters (location, amplitude) are used and their fields are summed to receive the field of the according tripole:

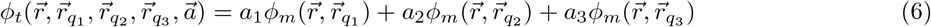

Accordingly

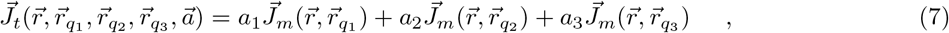

where *r_q_i__* is the location of source/sink *i* and *a_i_* the amplitude. In order to fulfill the necessity of closed current loops, the amplitudes *a_i_* have to sum to one, i.e. are in total normalized during modeling.

The tripolar model approaches tested within this paper are the following:

- *Tripole A:* One negative sink with amplitude –300 nA was placed at –1mm and one with –120 nA at 2.8 mm away from a central 420 nA source [Merletti and Farina, 2016].
- *Tripole B*: Two negative sinks (–210nA) were set –1.6mm and 3.2mm away from a central double positive source (420 nA) [Roeleveld et al., 1997].
- *Tripole C*: Two negative sources (–210nA) were set –2.1mm and 4.8mm away from a central double positive source (420 nA) [Kuiken et al., 2001].

As described above and shown in fig. 3, a tripoles scalp potential is not necessarily consisting of three peaks, even in proximity to the source. When using a tripole model with a non-equal distance between the monopoles such as the used *Tripoles A-C,* in particular one of the sources’ surface pattern peaks is weakened (the closer one). The resulting muscular *Tripoles A-C* are actually very similar to dipoles in the sense that the potential and current distributions are dominated by the stronger source-sink pair. This stronger source-sink pair was reached by either a larger distance between the sources or by larger source amplitudes.

## Deriving artefact model positions and orientations

For a realistic artefact source model, detailed muscle and eye positions within the head are essential. However, common MRI scans lack the necessary resolution and segmentation techniques to deduce the relevant anatomical details of muscles and eyes. Therefore, the open-source Multimodal Imaging-Based Detailed Anatomical (MIDA) model of the human head and neck [Iacono et al., 2015], an open-source, high-resolution MRI scan, segmented manually by experts, has been utilized.

### MIDA open-source Atlas

MIDA has been obtained by segmenting high-resolution T1- and T2-weighted MRI scans, magnetic resonance angiography (MRA), and diffusion tensor imaging (DTI) data of one healthy 29-year-old female volunteer. It was acquired at the Institute for Biomedical Engineering of the ETH Zurich (Switzerland) and contains 153 structures, segmented at 500 μm isotropic resolution.

### Surface mesh corrections

The segmented face and neck muscle meshes as depicted in the center of fig. 4 were corrected for selfintersections, unconnected nodes, and duplicated elements. Unconnected mesh parts were detected and removed. The triangulated surfaces were simplified, coarsened to an appropriate number of mesh vertices, and qualitatively enhanced (uniformity) while preserving the manifold.

**Figure 4:**
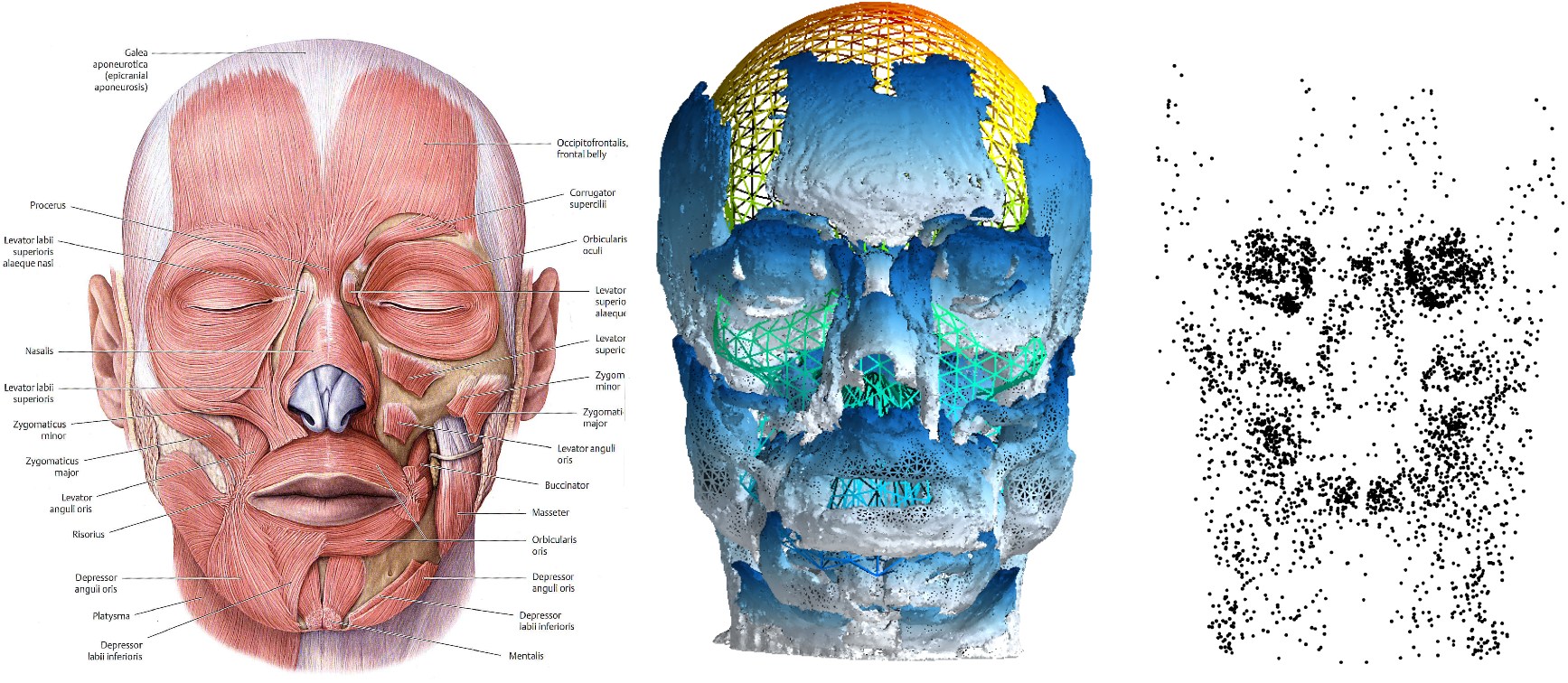
Comparison of Theory, MRI segmentation, and HArtMuTs artefact positions: Literature face muscle positions and directions (left, reprinted from Schünke et al. [2007], ©Thieme), muscular surface meshes from MIDA mapped onto the skull mesh of a BEM head model (middle) and HArtMuts final artefact source grid of size 3,9k (right).

### Artefact sourcemodel creation

Evenly spaced grids of 1 mm size were constructed within every muscle surface, building the artefact model’s source positions, of that a random sample of size 3,9k was drawn.^2^ According to the propagation direction of the MUAPs, the source’s pole orientations should follow the muscle’s fiber directions for simulation. The fiber directions were approximated by using a Principal Component Analysis (PCA) on close neighboring grid points of a muscles grid point cloud, where the closeness criteria have been varied depending on the underlying muscle shape.

For eye activity, source grids were built for six tissues in total. Firstly, the retina of the left and right eye was modeled with orientations pointing to the positive y-axis of the ACPC coordinate system (i.e. forward). Secondly, two source grids each were placed into the Cornea of the left and right eye, where one of them is oriented horizontally and the other vertically. To every position, a mirrored (on the x-axis) counterpart for the calculation of the symmetric dipoles was determined.

### Warping to MNI coordinates

A warping procedure for transferring the artefact sourcemodel from MIDA into MNI space was developed, which interpolates every point within the scalp mesh of one person into the scalp mesh of another person.

## Validation

HArtMuTs application to the inverse problem was validated with simulated EEG scalp patterns and subsequently on a real experimental EEG data set.

### Head model construction

A 4-shell Boundary Element Method (BEM) head model of the anatomy of Colin27 [Holmes et al., 1998] with neck extended scalp mesh was used for both validation steps. It consists of a cortex (gray + white matter), cerebrospinal fluid (CSF), skull, and scalp mesh and was segmented using an automatic segmentation pipeline^3^. After segmentation, the scalp mesh was merged with the neck part of the NYheads scalp mesh, in order to extend the scalp mesh of Colin27 by the neck part. The final surface meshes consist of 1922 vertices and 3990 triangles each. This has proven to be a reasonable compromise between model size, hence accuracy, and computational speed during inverse fit methods with inverse nonlinear fit requiring frequent leadfield recalculation [Miklody et al., 2016, Miklody, 2020]. As BEM solver OpenMEEG [Gramfort et al., 2010] was used, and the conductivity was fixed at the following values: scalp 0.465 S/m, skull 0.01 S/m, CSF 1.65 S/m, cortex 0.201 S/m.

### Inverse Fitting Routines

In order to make existing dipolar fitting routines work with the tripolar approach, we used the following approximation: Three tripoles were modeled around a central location in each direction of a Euclidean ACPC coordinate system. The linear combination of the three tripoles, that minimizes the error, was determined. This determined orientation and moment of the final single tripole and is the same procedure as for neural dipoles in particular with generic head models.

We used two different inverse fitting routines: *dipfit_gridsearch* and *dipfit_nonlinear* based on *FieldTrips*ft_dipolefitting [Oostendorp and Van Oosterom, 1989, Oostenveld et al., 2011]. Location and orientation of the sources were adjusted in an iterative procedure in an attempt to maximize the amount of variance in the data explained by the model. One criterion for evaluation and comparison of models, therefore, was the relative residual variance (RV), the relative amount of variance in the data, which is unexplained by the model [Scherg and Berg, 1991]. The RV is a sum-of-squares type error function, that computes the relative deviations of an estimated pattern 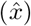 from the target pattern x:

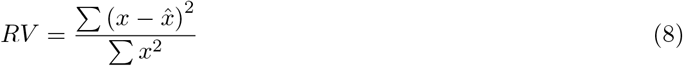

The RV has the advantage of being normalized to the average amplitude of the measured signal. *dipfit_gridsearch* compares the RV of the potentials at the electrodes (x) with the computed HArtMuT leadfields (x) for every source point in a gridded full sourcemodel (cortical + artefactual). For this gridded full sourcemodel, the sources were placed at a regular grid within the brain (cortical) and scalp (artefactual) meshes. Here, no restrictions were set to positions within the scalp, as e.g. from realistic positions derived from MIDA, in order to avoid systematic modeling errors due to incorrect segmentation or generic head models. The source position combined with an orientation, that produced the lowest RV with the scalp pattern, finally comprises the linear result to the inverse problem.

Subsequently, the nonlinear result for *dipfit_nonlinear* was achieved by allowing a minimization algorithm to vary orientation and source position using the *dipfit_gridsearch* results as starting points. In the end, a source is assigned the label of the closest artefact mesh for identification.

### Simulated Data

In the first validation step (simulated data), an FEM was used as a ‘ground-truth’ head model to validate the inverse fitting routines, the artefact positions with orientations and the 4-shell BEM HArtMut of Colin27 with extended neck as used for the inverse fit. HArtMuTs source localization results were compared to those of a standard 3-shell BEM model from EEGLAB [Delorme and Makeig, 2004a], which is also based on the anatomy of Colin27.

The simulated ‘ground-truth’ scalp patterns were calculated for all artefact positions with orientations from MIDA using only the common dipolar source model type, because no FEM solution for tripoles was available and the validity of the source localization is also given this way. As anatomy, the NYhead [Huang et al., 2016] was chosen, which is a high-resolution FEM in MNI space, that includes the neck. Its mesh and 75k cortical leadfields are publicly available. Since the published leadfields were calculated using the proprietary *Abaqus FEA software suite,* the cortical leadfields were recalculated using the open-source software package SimBio [SimBio Development Group, 2022]. Comparing both the original Abaqus Software and the SimBio leadfield simulations, an average correlation of 0.947 for single cortex source location leadfields was found. The discrepancy may be explained by the use of the different solvers and an additional linear transformation of source points and electrodes applied beforehand. This transformation was needed to bring the published FEM mesh and source and electrode positions into the same coordinate system.

Using the NYhead, FEM leadfields of a 3540 large artefact source model from MIDA were calculated at the same 129 aligned electrodes as in the dataset used in the experimental data validation. Each leadfield was considered as a single scalp pattern, to which the corresponding source position had to be localized.

### Results of the validation on simulated data

The results of the source localization on simulated dipolar cortical and dipolar artefactual source points are plotted in fig. 5. The standard EEGLAB model led to significantly higher RV and source localization error for both brain and non-brain sources than HArtMuT dipoles *(p* ≤ 0.005). The average source localization error for brain and artefact sources using HArtMuT dipoles was estimated to be below 10mm with an RV mainly below 0.01. The median RV was 0.0032 for the brain and 0.0042 for artefacts. The median source localization error was 8.4 mm for the brain and 9.6 mm for artefacts. In comparison, the standard EEGLAB model has a similar median source localization error of 9.1 mm for the brain, but a much higher value of 34.2 mm for non-brain sources. The median RVs were 0.010 for brain and 0.25 for non-brain sources.

**Figure 5:**
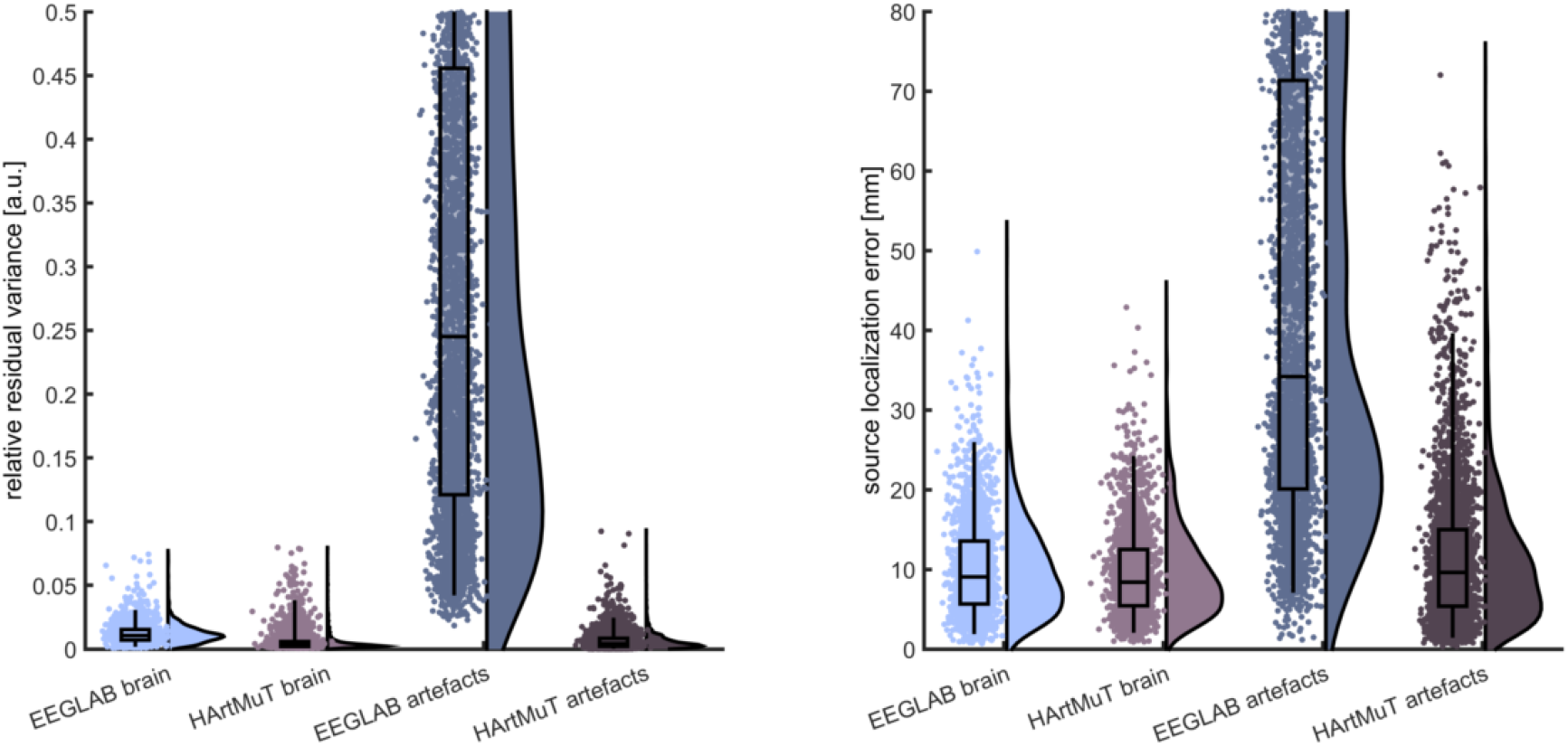
Source localization results of HArtMuT dipoles and the EEGLAB model compared for simulated FEM data. The relative residual variance (RV) and the geometrical error in source localization using *dipfit_nonlinear* for FEM-simulated scalp potentials reconstructed using the 4-shell HArtMuT dipoles remained reasonably low (RV median 0.025 and source reconstruction error around 10mm). The geometries of the two models (NYhead vs. Colin27) were not exactly the same, such that the electrode and source positions needed to be transformed, mimicking a realistic scenario.

### Remarks on the validation with simulated data

The validation against simulated data from a HD-FEM model as ‘ground-truth’ was successful. This confirmed in particular HArtMuTs 4-shell BEM model as a whole, the used inverse fitting routines, the neck extension procedure developed and applied to Colin27, and furthermore the artefact translation procedure between the different head shapes (MIDA, NYhead, Colin27).

## Experimental data

The second validation step (experimental data) was performed by using data from an EEG experiment [Gramann et al., 2021]. From the experimental data, different components and their scalp patterns were extracted using ICA within EEGLAB [Delorme and Makeig, 2004b]. During source localization using HArtMuT, the performance of the different artefact modeling types (dipole, tripoles, symmetric dipole) was evaluated. The RVs were compared to see, which source model type best describes the experimental data patterns, and thus, which source model type overall allows for the best fit for the different types of measured signals in EEG. For comparison, the source localization results of the commonly used standard EEGLab model were computed as well.

### Data and processing

The data of 19 healthy subjects (11 female, 8 male, aged 20-46 years, 30.25 years on average) were recorded in a Mobile Brain/Body Imaging study [Gramann et al., 2021], focusing on heading computation in a stationary and a mobile experimental condition. The virtual environment consisted of a simple floor rendering without external landmarks. Each trial started with a pole (indicating zero heading) which was then replaced by a sphere moving around the participant at a constant distance but with different velocity profiles. Participants were instructed to follow the movement of the sphere and to rotate back after the sphere stopped moving unpredictably at eccentricities between 30 and 150 degrees to the left or right. In the stationary condition, participants followed the sphere using a joystick while standing in front of a 2D monitor. In the mobile condition, they physically rotated wearing a virtual reality head-mounted display. Rotation eccentricity, speed, and direction were varied across 140 trials per movement condition (stationary vs. mobile). For the present study and modeling approaches, only the full-body rotation condition was taken into account, as we were interested in the contributions of neck muscles and eye movements to the sensor signal.

EEG data were recorded with a 1 kHz sampling rate from 157 active electrodes (BrainProducts, Gilching, Germany) with 129 channels in a cap with an equidistant layout (with two vertical EOG channels, both eyes) and 28 channels in a custom neckband. Individual electrode locations were recorded (Polaris Vicra, NDI, Waterloo, ON, Canada). Additional motion data was recorded but is not considered for the modeling approach in this study.

The data was processed in MATLAB (R2016b version 9.1; The MathWorks Inc., Natick, Massachusetts, USA) using custom scripts based on the EEGLAB toolbox [Delorme and Makeig, 2004a, version 14.1.0]. The 28 channels located in the neckband did not increase the decomposition quality of the ICA [Klug and Gramann, 2021] and were thus removed. Subsequently, the data were re-sampled to 250 Hz, line noise was removed with Zapline [de Cheveigné, 2020] using default settings, and breaks and pre-/post-experiment segments were removed from the data. We then used the *clean_rawdata* function of EEGLAB to detect and interpolate bad channels (disregarding EOG channels) using an 0.8 correlation threshold and no other measures. Subsequently, the data were re-referenced to the average of all channels except the EOG channels. We then applied a 1 Hz high-pass filter as suggested by Klug and Gramann [2021] and computed an ICA using the AMICA algorithm [Palmer et al., 2011] with 2000 iterations and automatic rejection of samples (3 times with 3 SD threshold).

In summarizing, the 19 subjects had on average 112 ICs depending on the number of valid channels after the preceding artefact channel rejection and rank of the data. In total 2132 ICs were extracted and to each IC a source position had to be localized using the above described dipfit routines from EEGLAB. Finally, the components were labelled using the ICLabel ‘lite’ classifier [Pion-Tonachini et al., 2019].

### Results

Of the total 2132 extracted components of all subjects, 69.4% were detected as muscle sources based on HArtMuTs source location, while 28.8% were brain and around 1.7% eye sources. The amplitudes of eye sources were overall strongest (median 8.8 *μ*V, maximum 419.0 *μ*V) followed by muscle sources (median 0.4*μ*V, maximum 181.1 *μ*V) and brain sources (median 0.2*μ*V, maximum 36.0*μ*V).

Comparing different approaches for modeling the muscles in fig. 6, *Tripole B* led to the lowest median and lower quartile errors, which is why it was also used as the main tripolar model of HArtMuT tripoles, although the differences are not significant.

**Figure 6:**
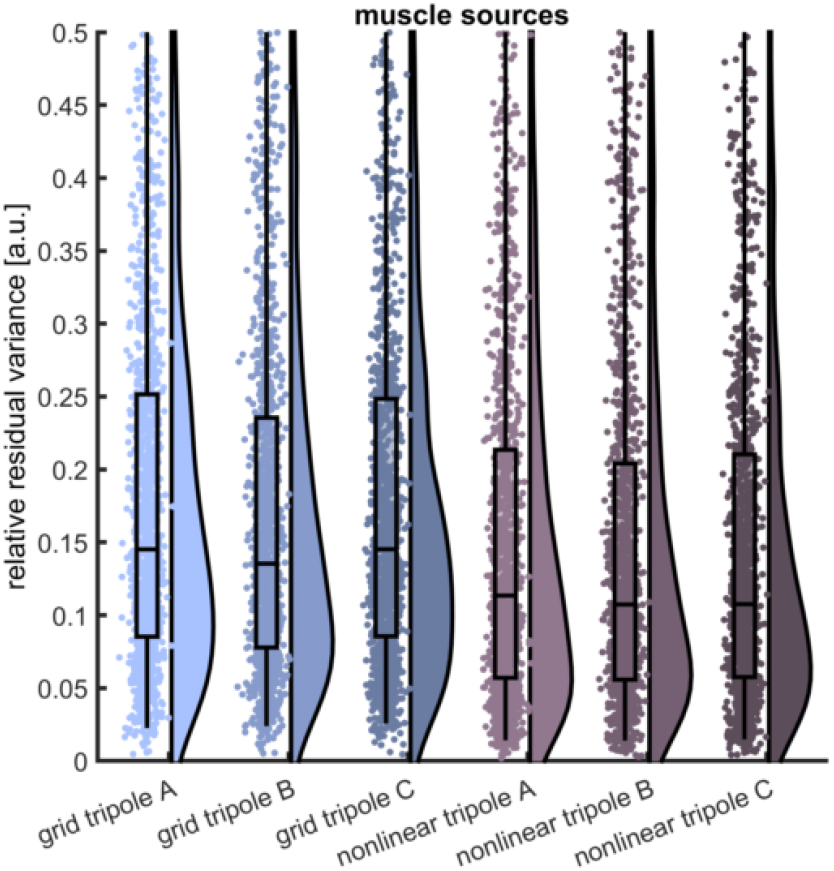
The residual variance of the muscular source reconstruction results for the three *Tripoles A-C* using *dipfit_gridsearch* (grid) and *dipfit_nonlinear* (nonlinear) as different source localization algorithms: *Tripole B* led to lowest, *Tripole C* to second lowest, and *Tripole A* to highest errors. The differences were not significant.

Most of the muscular IC patterns were localized close to the combination of Muscles temporalis and temporoparietalis (21.6%), followed by MIDAs unspecific Muscle (M.) generalis class (18.1%), which is a collection of muscular tissue in the lower neck. Then it follows the M. splenius capitis (4.2%) and M. Sternocleidomastoid (4.1%). All other muscles (26 in total) were below 4% in their occurrences.

For the eyes, using dipolar models that are symmetrically aligned (mirrored at the plane spanned by the x- and z-axis in the ACPC coordinate system) with equal amplitudes led to the lowest error using *dipfit_gridsearch* compared to any other combination (fig. 7). Since fitting symmetric dipoles with *dipfit_nonlinear* was not implemented, the final results of the symmetric dipole fit remain those from *dipfit_gridsearch*.

**Figure 7:**
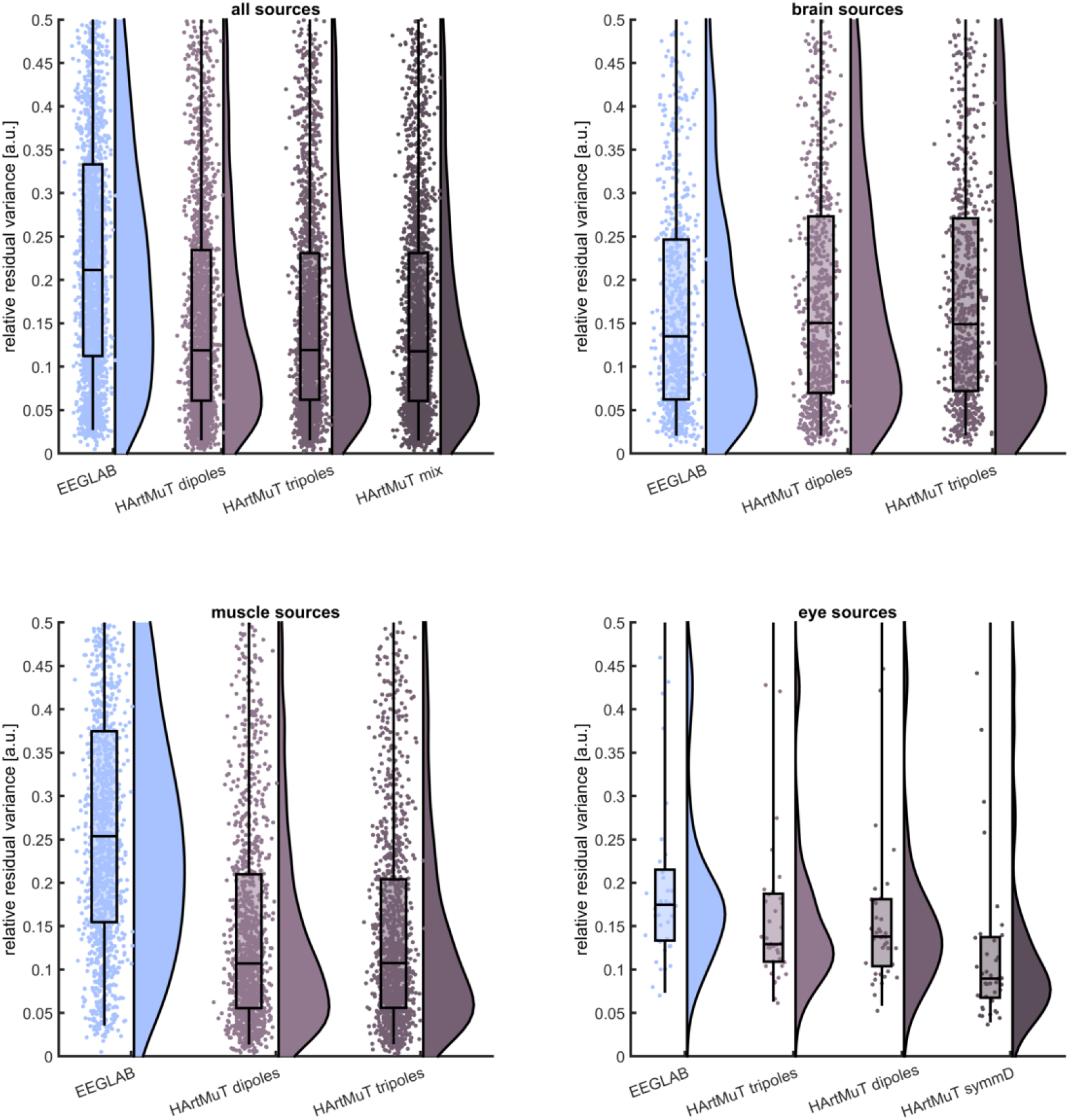
The relative residual variance (RV) of all (upper left), cortical (upper right), muscular (lower left) and ocular (lower right) source reconstruction results for different source models using *dipfit_nonlinear* over 19 subjects: HArtMuT mix (dipoles for the brain, tripoles for muscle, and symmetric dipoles for eye sources) led to the lowest errors overall.

For further analysis, the best models for each type of source, namely cortical dipoles, muscular tripoles (*Tripole B*), and ocular symmetric dipoles, were combined to one model, which will be referred to as HArtMut mix in the following.

For cortical sources, dipoles led to significantly *(p* ≤ 0.001) lower errors than tripoles (1/19) while the standard EEGLAB model produced the lowest errors (19/19). All types of HArtMuT (dipole, tripole, symmetric dipole) lead to significantly lower errors for all subjects (p ≤ 0.05) for muscles than the standard EEGLAB model. Tripoles lead to significantly lower errors for muscles than dipoles in 2/19 subjects (*p* ≤ 0.05) and for two additional participants very close to significance (*p* ≤ 0.05). Symmetric dipoles (symmD) led to the lowest errors for eye sources while significant only for 2/19 subjects most probably due to the low number of sources per subject (0-5, 2 on average). The standard EEGLAB model significantly (*p* ≤ 0.05) produced the highest RVs in all subjects for muscle and eye sources. Note, that the symmetric dipoles (symmD) were only fitted using *dipfit_gridsearch* and could potentially be further improved by *dipfit_nonlinear*.

The performance of HArtMuT mix is demonstrated in fig. 7, which compares the results of HArtMuT with those of the standard EEGLAB model for brain, muscle, and eye sources separately. For cortical sources, the standard EEGLAB model produced slightly but still significantly lower errors than HArtMuT (*α* ≤ 0.01). Using HArtMut mix, the RVs were reduced from a median 0.25 to 0.11 for muscle sources and 0.18 to 0.09 for eye sources compared to using the standard EEGLAB 3-shell model. With the simple *dipfit_gridsearch,* HArtMuT tripoles reduced the RV compared to HArtMuT dipoles for 3/19 subjects (*α* ≤ 0.05) only. When further optimized by *dipfit_nonlinear,* the differences in RV between HArtMuT dipoles and HArtMuT tripoles were not significant anymore except for one subject (*α* ≤ 0.05). Using *dipfit_nonlinear* significantly (*α* ≤ 0.05) reduced the errors within any source model in every subject.

### Labeling

We also investigated the resulting classification based on source location with automatic IC classification using IClabel, an EEGLAB plugin. IClabel automatically classifies IC components into different classes based on a classifier, which was built with a crowd labeling approach [Pion-Tonachini et al., 2019]. The comparison of IClabel and HArtMuTs classification is summarized in tab. 1.

**Table 1:** Correspondence between IC classification based on IClabel and localization in the HArtMuT mix. Of the 15.6% brain sources labelled by IClabel, 12.9% were also found as brain sources by HArtMuT mix, which corresponds to an overlap of 83% of all brain sources. Of the 42.4% muscles even 39.6% were detected also by HArtMuT mix as muscle sources, such that the match is 93%. For the eyes, however, the 1.7% of all 3.0% that both labelled as eyes, correspond to only 40% match. IClabels line and channel noise, heart and other labels sum to in total 38.9% of all ICs, to which HArtMuT mix assigned mostly muscle (23.7%) or brain (14.8%) sources.

|  |  | IClable |  |  |  |  |  |  | sum |
| --- | --- | --- | --- | --- | --- | --- | --- | --- | --- |
|  |  | brain | muscle | eye | heart | line noise | channel noise | other |  |
| **HArtMuT mix** | brain | 12.9% | 2.3% | 0.1% | 0.0% | 0.8% | 0.4% | 13.6% | **30.1%** |
|  | muscle | 2.7% | 39.6% | 1.7% | 0.3% | 0.6% | 6.8% | 16.0% | **67.7%** |
|  | eye | 0.0% | 0.5% | 1.2% | 0.0% | 0.0% | 0.0% | 0.4% | **2.1%** |
|  | **sum** | **15.6%** | **42.4%** | **3.0%** | **0.3%** | **1.4%** | **7.2%** | **30.0%** |  |

This simple approach, in which only the comparison of the scalp patterns of the forward model with IC patterns is incorporated, already leads to a high correspondence between the IC classification labels and the resulting source location based classification. Of the ICs classified as brain by IClabel, there were 83% equally classified as brain by HArtMuT mix and for IClabels muscles 93% as muscles by HArtMuT mix. The median RV of IC classification as ‘other’ was 0.22 opposing all remaining classes’ RV of 0.096. The components classified as ‘eye’ by IClabel were 40% classified as eyes and 57% as muscles. It is noteworthy, that 0.9% of the by HArtMuT mix localized muscle sources were actually eye muscles and additionally 2,7% eyelid muscles (Muscle Orbicularis Oculi), which one might also label as eye artefact.

This large correspondence between HArtMuT and IClabel is remarkable, although one has to take into account, that it is not assured that IClabel classified correctly.

### Comparison with physiologically motivated artefact sourcemodel

Of the 1517 reconstructed artefact sources, only 324 (21.4%) were found to be actually inside an artefact mesh as derived from MIDAs MRI segmentation. However, the median distance of a source to the artefact mesh corresponding to its label was 4.5 mm with a standard deviation of 52.7 mm. The relatively high standard deviation is caused by a few outliers, such that 74% (54%) are closer than 10 mm (5 mm) to its mesh. Interestingly, these outliers have in most cases high RVs, meaning that they stem from rather unsuccessful fits. If one for example restricts this analysis to poor RVs, e.g. > 0.3, which corresponds to 20% of the data, the mean distance of a source to the artefact mesh corresponding to its label is 35.7mm (84.8 mm standard deviation).

### Detailed view on muscular and ocular source model type performances

Looking at the exemplary source localization results and the corresponding patterns reveals several insights. First of all, the scalp patterns of muscular sources appeared as expected as common muscular artefact patterns and tripoles almost always led to lower RVs. However, it was surprising to us, that a dipole can lead to a tripolar-like pattern in the EEG like that measured in the upper row of fig. 8. Investigations revealed that this was partly because of interpolation artefacts and partly because sources normally oriented and very close to the scalp surface can actually lead to similar patterns in particular in regions with heterogeneous scalp and skull thickness like the forehead. However, in most cases, a well-fitted tripole still remained the better model here. The lower row of fig. 8 shows another example from the back of the head in the neck region with a slightly higher RV. Here, both dipole and tripole do not fully reproduce the shape of the positive peak but the approximation is still reasonable. What can be seen in both examples of fig. 8 is, that some of the seemingly multi-poled solutions are rather dipolar but interpolation of the EEG scalp plotting from EEGLAB led to very strong (in this case negative) peaks in areas where there were no electrodes (no black dots). This means, that a comparison of BEM solver solutions in form of scalp plots, might not always be a good evaluation approach.

**Figure 8:**
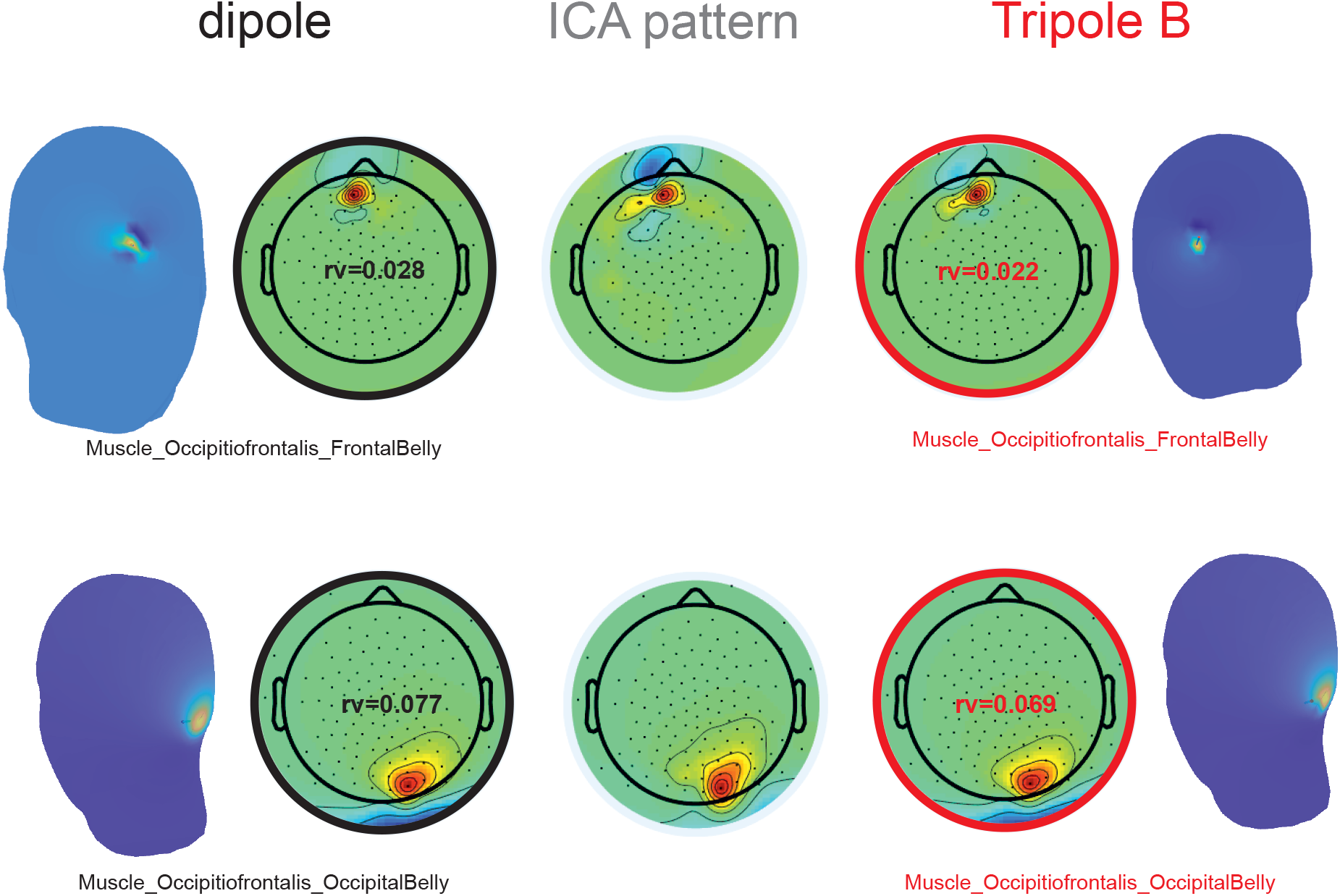
Two exemplary source models and their solution in source localization for muscle artefacts: the center shows the IC extracted from the data, in the left is the dipolar, and in the right the tripolar source on the scalp and their corresponding EEG patterns. Black is the dipolar model and moment, red tripolar with Tripole B.

In fig. 9 the clear difference between symmetric and non-symmetric dipolar eye models can be investigated. As we expected, the best eye models concerning the pattern reconstruction error were linked symmetrically. Here, the simple *dipfit_gridsearch* outperforms all other models that use non-symmetric tripoles and dipoles. In the source locations we see that for these models the source was mostly located somewhere between the eyes, which is not realistic.

**Figure 9:**
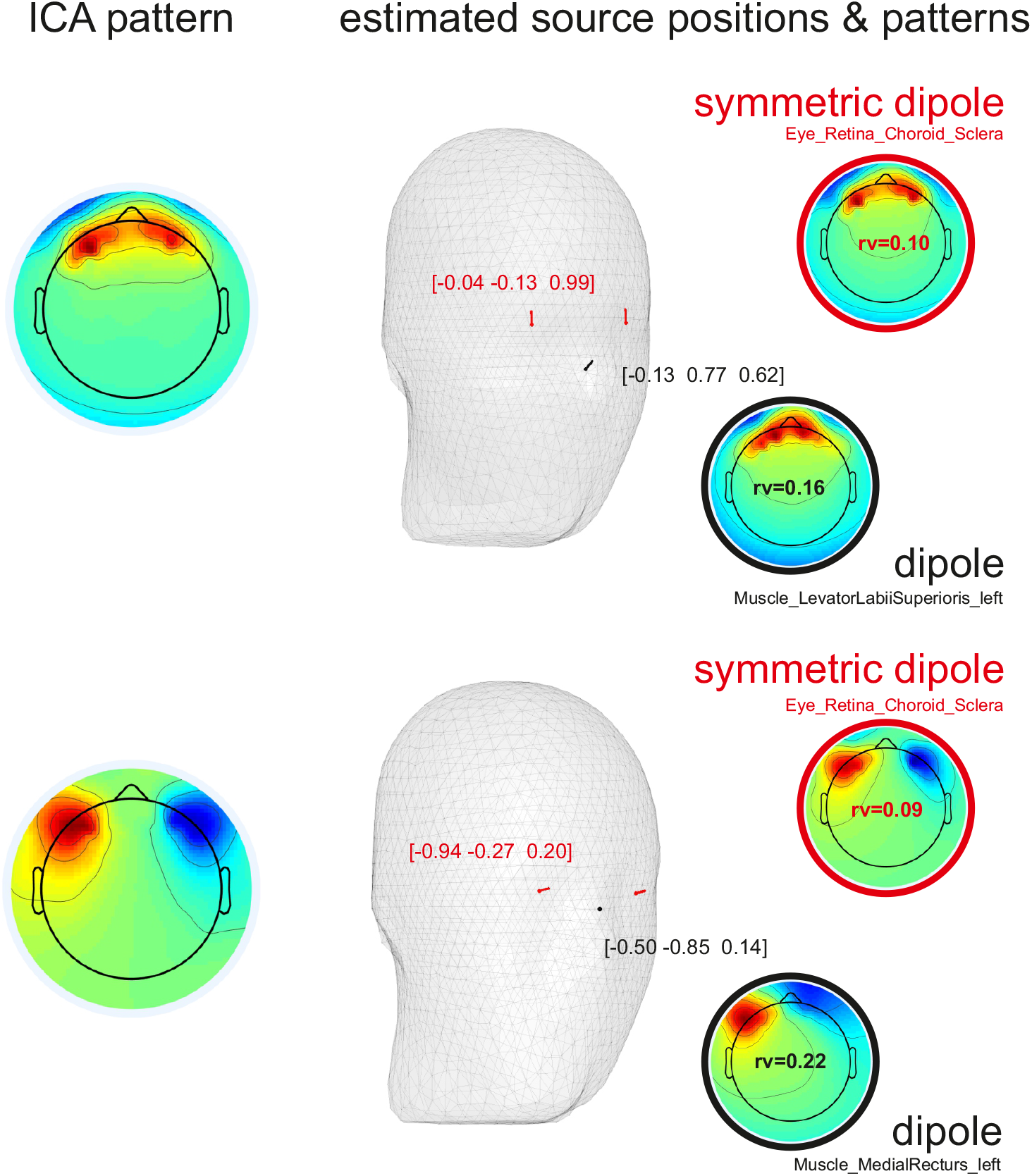
Comparison of symmetric dipolar (red) and dipolar (black) source models and their solution in source localization on two exemplary eye artefacts. On the left, one can find the IC extracted from the data, in the center is the fitted location and moment in the head, while on the right you find their corresponding patterns, RVs, and labels. The upper row shows an example IC of vertical eye movement, whereas in the lower row an exemplary IC of horizontal eye movement is shown. In both examples, the superiority of the symmetric dipole over the simple dipole is visible.

Most ocular sources were found to lie in or close to the Retina (84%), while the rest lie in or close to the Cornea (16%). Interestingly, the commonly found pattern on the bottom in fig. 9 corresponds to the straight forward looking eye position. The approach only catches static eye positions in single ICs and eye movements will be spread over linear combinations of static patterns such as the upper for upwards movements away from the lower straight forward position. A *dipfit-nonlinear* routine for linked symmetric positions had not been implemented within this work but is expected to further improve the results.

## Discussion

We investigated new approaches to modeling muscle and eye contributions to EEG and used the models in source reconstruction on simulated and experimental data.

In sum, we propose our new head model HArtMuT, which includes the neck and corresponding brain, muscle, and eye tissue source positions and labels. It is available for different head anatomies and BEM/FEM models. We showed the possibility of including artefacts into common source localization routines and validated HArtMuT on simulated data. We tested the effect of modeling sources as dipoles, tripoles, and symmetric dipoles and measured the performance on experimental data in residual variance. While a single dipole model is most likely sufficient for reasonably good source localization of all components except the eyes, it is more precise to use a tripolar model for muscle sources. This is in accordance with the triphasic MUAP and did slightly improve the estimation of source potentials for muscles for some of the subjects, but differences were not significant. If tripoles are not at hand, a dipolar model, which is commonly implemented in FEM/BEM solvers, can also be physiologically justified, as the triphasic MUAP shows a strong dipolar behavior followed by a weaker AHP bump.

For the eyes, a symmetric dipolar model led to the lowest approximation errors, which reflects the synchronous movements of the eyes. This met our expectations since ICA decompositions cannot differentiate the signals of both eyeballs.

To summarize, the best option is a mixture of the models depending on the source’s tissue type. This best combination consists of dipoles for the brain, tripoles (*Tripole B*) for the muscles, and symmetric dipoles for the eyes. This is condensed into one model: HArtMuT mix.

Since the used tripolar models *Tripoles A-C* were all motivated by literature, there is the potential to further improve the tripolar sources localization results by improved architectures. Underlining this, we achieved promising preliminary results by optimizing positions and amplitudes of three single monopolar sources freely in the head by non-linear optimization. However, the success drastically depends on the initial starting point (we used the tripolar solution) and the optimization gets easily stuck at local minima. Such solutions however often lead to lower RVs and more convincing scalp patterns. We plan to further elaborate on this approach.

Not covered in this article, we also investigated extending the scalp mesh by the upper torso to for example include the possibility of localizing signals of the heart. However, neither any meaningful source had been localized in that region nor did the RV improve. This also indicates that the chosen neck extension of HArtMuT covers neck muscular activity in real EEG data sufficiently.

It remains an open question whether the case of *dipfit_nonlinear* optimized 3-shell RV of brain ICs being significantly below the other for all subjects means that our model is not optimal here, or that the 3-shell just produces lower RVs that do not directly correspond to the actual source localization error. In general, the RV tells only about fitting quality from a numeric perspective, not plausibility. The most direct interpretation of this is that the 3-shell is the better-fitted model for the purpose of source localization. The 4-shell is more detailed but delivers more systematic errors either concerning the geometry or in combination with a not fully appropriate source model (dipoles). In contrast, a thorough post-hoc investigation of sources from the brain revealed that many of the ICA patterns were either a mixture of multiple dipolar sources or scalp plots suggesting that a distributed source model could make more sense. In both cases, the solutions of the EEGlab standard model were mostly the better fit due to its higher spatial spread of the single sources (more spatial smearing). Additionally, the results of the validation on an FEM model proved that the 4-shell model is the better model also for brain sources and we hence believe that the main reason for the 3-shell performance on experimental data was rather a non-optimal IC decomposition than head model.

Another factor might be non-optimal tissue conductivities in the 4-shell head model. A post-hoc analysis with preliminary results revealed that HArtMuTs RVs can be improved for the brain to a similar level by fitted conductivities. A further reason could be the missing anatomical correspondence with the subjects. One might conjecture that the 3-shell is delivering lower errors because it is more generic with its constant skull thickness. However, more investigation on this is needed.

A similar argumentation can be brought up to explain the relatively large source localization error for both models HArtMuT and standard EEGLAB in the first validation step (using the FEM leadfields as ground truth). Besides not matching conductivities, in particular different solvers (FEM vs. BEM), different anatomies (NYhead vs. Colin27), and needed transformations of electrode and source positions are to name here as origins of uncertainties, that summed up in the end can lead to such an average source localization error of approx. 9 mm.

All together, HArtMuT significantly improved source localization compared to the commonly used EEGLAB model in particular for muscle and eye sources. Furthermore, muscle and eye artefacts could clearly be differentiated from brain sources by their patterns via inverse fitting routines together with an appropriate model like HArtMuT. A comparison with IClabel indicates that HArtMuT may be a promising tool not only for source localization but also for classification of ICs.

It is however noteworthy, that IClabel classifications cannot be seen as the ‘ground truth’ in all studies, because it was mostly trained on EEG data from stationary participants. In this study, IClabels classifications were introduced to serve a basic intuition for the quality of HArtMuTs results. A further study is planned to investigate the usage of HArtMuT for artefact labels in detail.

## Supplementary Material

The final HArtMuT (BEM- and FEM-models) including cortical and artefactual sourcemodels, leadfields and labels are publicly available^4^.

## Acknowledgements

We want to thank Benjamin Blankertz for his outstanding general and logistical support of this project. N.H. and K.G. acknowledge support from the Deutsche Forschungsgemeinschaft (DFG, German Research Foundation): N.H. from GRK 2433/1, Project number 384950143, and K.G. from GR 2627/8-1. Open access funding enabled and organized by Projekt DEAL.

## Contributions

N.H. and D.M. implemented the segmentation and warping routines. N.H. deduced the artefact atlas for head models, researched mathematical approximation and physiological origin of artefacts and implemented the FEM head model. D.M. took care of the BEM head model including tripolar model implementation. M.K. implemented the first versions of the dipfit routines in EEGLAB, which were then further improved. K.G. had the original idea and was closely supporting the project in the beginning and in the final manuscript phase. All authors contributed to writing of the manuscript.

## Footnotes

1 https://github.com/harmening/HArtMuT

2 For the source reconstruction, however, regularly spaced grids were used as described below.

3 https://github.com/harmening/MRIsegmentation

4 https://github.com/harmening/HArtMuT

